# FIGL1 and its novel partner FLIP form a conserved complex that regulates homologous recombination

**DOI:** 10.1101/159657

**Authors:** Joiselle Blanche Fernandes, Marine Duhamel, Mathilde Séguéla-Arnaud, Nicole Froger, Chloé Girard, Sandrine Choinard, Nancy De Winne, Geert De Jaeger, Kris Gevaert, Raphael Guerois, Rajeev Kumar, Raphael Mercier

## Abstract

Homologous recombination is central to repair DNA double-strand breaks (DSB), either accidently arising in mitotic cells or in a programed manner at meiosis. Crossovers resulting from the repair of meiotic breaks are essential for proper chromosome segregation and increase genetic diversity of the progeny. However, mechanisms regulating CO formation remain elusive. Here, we identified through protein-protein interaction and genetic screens FIDGETIN-LIKE-1 INTERACTING PROTEIN (FLIP) as a new partner of the previously characterized anti-crossover factor FIDGETIN-LIKE-1 (FIGL1) in *Arabidopsis thaliana*. We showed that FLIP limits meiotic crossover together with FIGL1. Further, FLIP and FIGL1 form a protein complex conserved from Arabidopsis to Human. FIGL1 interacts with the recombinases RAD51 and DMC1, the enzymes that catalyze the DNA stand exchange step of homologous recombination. Arabidopsis *flip* mutants recapitulates the *figl1* phenotype, with enhanced meiotic recombination associated with change in DMC1 dynamics. Our data thus suggest that FLIP and FIGL1 form a conserved complex that regulates the crucial step of strand invasion in homologous recombination.

---

Homologous recombination (HR) is critical for the repair of DNA double-strand breaks (DSBs) in both mitotic and meiotic cells^1^. Defects in HR repair causes genomic instability, leading to cancer predisposition and various inherited diseases in Humans ^2^. During meiosis, HR promotes reciprocal exchange of genetic material between the homologous chromosomes by forming crossovers (COs). COs between the homologs constitute a physical link which is crucial for the accurate segregation of homologous chromosomes during meiosis ^3^. COs also reshuffle parental genomes to enhance genetic diversity on which selection can act ^4^. Failure or errors in HR at meiosis leads to sterility and aneuploidy, such as Down syndrome in humans ^5,6^.

During meiosis, HR is initiated by the formation of numerous programmed DSBs catalyzed by the topoisomerase-like protein SPO11^7^. DSBs are resected to form 3’ single-stranded DNA (ssDNA) overhangs. A central step of HR is the search and invasion of an intact homologous template by the broken DNA end, which is catalyzed by two recombinases, RAD51 and its meiosis-specific paralog DMC1 ^8^. Both recombinases polymerize on 3’ ssDNA overhangs to form nucleoprotein filaments that can be cytologically observed as foci on chromosomes ^9,10^. At this step, meiotic DSB repair encounters two possibilities to repair DSB by HR, either using the sister chromatid (inter-sister recombination) or using the homologous chromosomes (inter-homolog recombination).

The invasion and strand exchange of ssDNA displaces one strand of the template DNA, resulting in a three-stranded joint molecule (D-loops). D-loops are precursors for different pathways leading to either reciprocal exchange (CO) or non-reciprocal exchange (NCO) between the homologous chromosomes. Two pathways of COs formation, classified as class I and class II, have been characterized, with variably relative importance in different species^3^. Class I COs are dependent on the activity of a group of protein collectively called ZMM (for Zip1-4, Msh4-5, Mer3)^11^, which stabilize D-loop intermediates to promote formation of the double-Holliday junction intermediates^12^. MLH1 and MLH3 in conjunction with EXO1 resolve double-Holliday junctions as class I COs ^13,14^. Class I CO occurrence reduces the probability of another CO forming in the vicinity, a phenomenon termed as CO interference ^15^. Additionally, recombination intermediates can be resolved by structure specific endonucleases including MUS81, producing class II COs, which are not subjected to interference ^16–18^. In Arabidopsis, class I COs constitute 85-90% of COs, while remaining minority are class II CO ^19,20^. Like in most eukaryotes, DSBs largely outnumber COs in Arabidopsis with a ratio of ~25:1 ^21^. This suggests that active mechanisms prevent DSBs from becoming CO. Accordingly, several anti-CO factors are identified in different species ^10,22–30^.

Previously, our forward genetic screen identified FIDGETIN-LIKE-1 (FIGL1) as a negative regulator of meiotic COs in Arabidopsis ^10^. Mutation in *FIGL1* in *Arabidopsis* increases meiotic CO frequency by 1.8-fold compared to wild type, and modifies the number and/or dynamics of RAD51/DMC1 foci. FIGL1 is widely conserved and is required for efficient HR in human somatic cells through a direct interaction with RAD51 ^31^. Altogether, this suggests that FIGL1 is a conserved regulator of the strand invasion step of recombination, both in somatic and meiotic cells. FIGL1 belongs to the large family of AAA-ATPase proteins that are implicated in structural remodeling, unfolding and disassembly of proteins and oligomer complexes ^32,33^.

Here, we identified a new factor limiting COs in Arabidopsis and that interacts directly with FIGL1, which we named FIDGETIN-LIKE-1 INTERACTING PROTEIN (FLIP). FLIP and its interaction with FIGL1 are conserved from plants to mammals, which suggests that the complex was present at the root of the eukaryotic tree. We further showed that *FLIP* and *FIGL1* act in the same pathway to negatively regulate meiotic CO formation, which appears to act on the regulation of the recombinase DMC1. Finally, we showed that both Arabidopsis and human FIGL1/FLIP complex interact with both RAD51 and DMC1. Overall, this study identified a novel conserved protein complex that regulates a crucial step of homologous recombination.

## RESULTS

### Identification of FIDGETIN-LIKE-1 Interacting Protein (FLIP), an evolutionarily conserved partner of FIGL1

We previously identified FIDGETIN-LIKE-1 (FIGL1) as an anti-CO protein^10^. To better understand the role of FIGL1 during meiotic recombination, we searched for its interacting partners by tandem affinity purification coupled to mass spectrometry (TAP-MS) using overexpressed FIGL1 as a bait in *Arabidopsis* suspension culture cells ^34^. (Figure 1A). After filtering co-purified proteins for false positive (see M&M and ^34^), we recovered, in two independent experiments, peptides from a single protein encoded by a gene of unknown function *(AT1G04650),* and therefore named it as FIDGETIN-LIKE-1 INTERACTING PROTEIN (FLIP). Reciprocal TAP-MS experiments using FLIP as bait also recovered FIGL1 peptides, further suggesting that FLIP and FIGL1 belong to the same complex *in vivo* (Figure 1B). A direct interaction between FLIP and FIGL1 was further supported by yeast two hybrid (Y2H) assay using full length proteins (Figure 1C). To map the interaction domains, we truncated FIGL1 and FLIP proteins and tested their interaction in Y2H assays. N-terminal regions of FLIP (1-502 aminoacids) and of FIGL1 (1-271 aminoacids), lacking the FRBD (FIGNL1’s RAD51 binding domain) and the AAA-ATPase domain, were sufficient to mediate their interaction (Figure 1C). Further, the N terminal domain of FLIP was able to interact with itself, suggesting that it could oligomerize. Moreover, the human ortholog of FLIP (C1ORF112, hFLIP) and FIGL1 (hFIGNL1) also showed interaction in our Y2H assays, suggesting that this interaction is evolutionarily conserved (Figure 1D). In addition, hFIGNL1 and C1ORF112 proteins were previously showed to co-purify in pull-down assays^31,35^. and mouse corresponding genes are strongly co-expressed ^36^, further supporting that the FIGL1-FLIP interaction is conserved from plants to mammals. Mapping of the interaction domain on human proteins showed the N-terminal region (1-290 aminoacids) of FIGNL1 to mediate the interaction with hFLIP, consistent with the Arabidopsis data (Figure 1C and 1D). In addition, the FRBD of hFIGNL1 shows an interaction with hFLIP, suggesting that the FRBD domain could also participate to the interaction.

**Figure 1:**
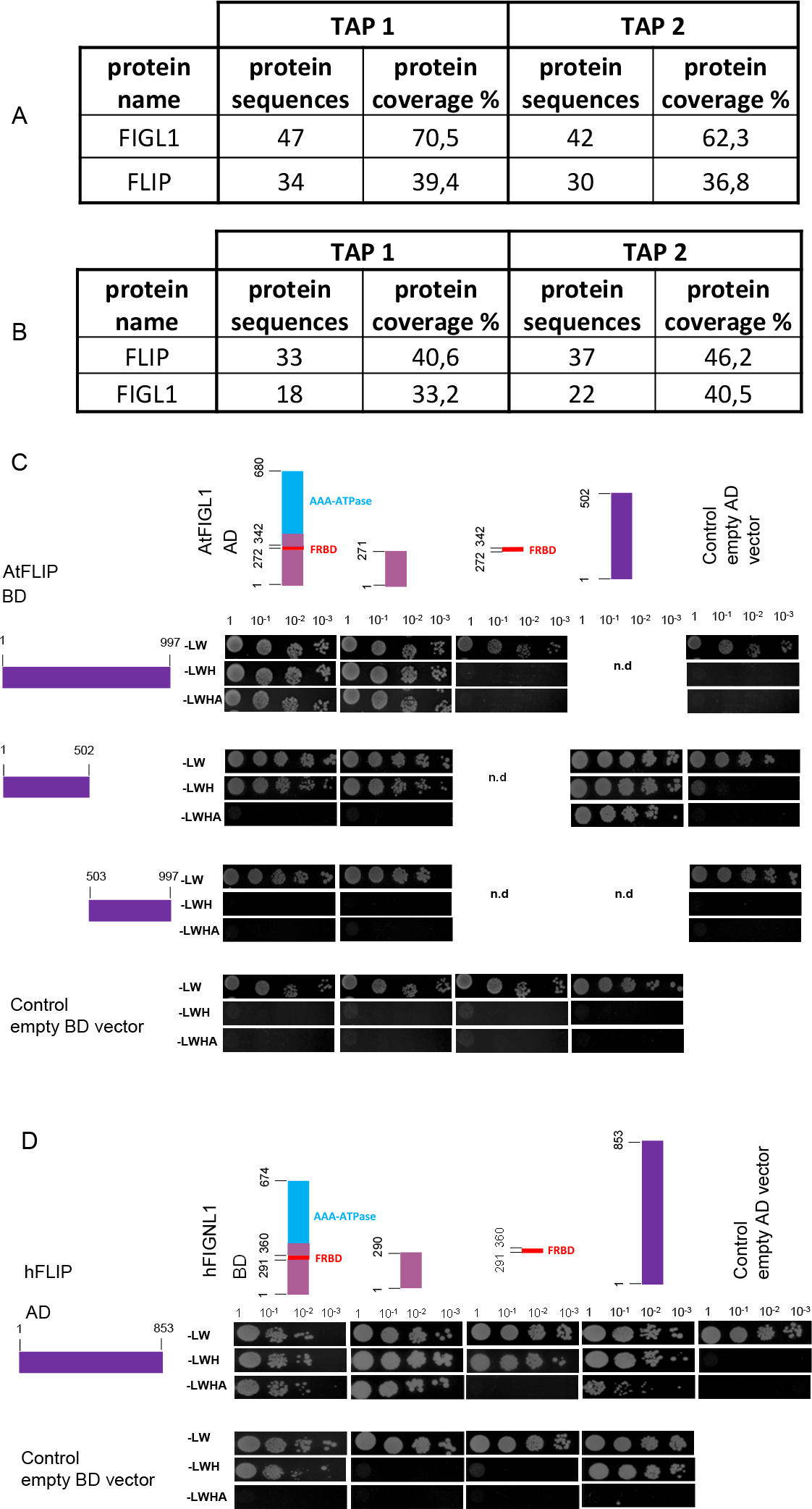
*Arabidopsis* and Humans FLIP and FIGL1 interact with each other through N-terminal. A-B. Represents purified proteins in two replicates of tandem affinity purifications followed by mass spectrometry, performed using *Arabidopsis* FIGL1 (A) or FLIP (B) as a bait, over-expressed in *Arabidopsis* cultured cells. For filtering aspecific and false positive interactors, we refer to M&M and ^34^. The number of peptides and the fraction of the protein covered are indicated for each hit. Detailed information is present in Table S1. C. *Arabidopsis* FLIP-FIGL1 interact using yeast two hybrid assay. Schematic representation of full length and truncated proteins. Position of the truncation and the known domain are indicated. The two protein of interest are fused with GAL4 DNA binding domain (BD) and with GAL4 activation domain (AD), respectively. Diploid strains are selected on medium lacking LW amino acids. Protein interactions were observed by selecting diploids grown on medium stringency media lacking LWH or in high stringency medium lacking LWHA. Controls empty AD and BD vector are shown. Serial dilutions are indicated in the order 1, 10, 100, 1000. Combinations of fusion proteins not tested are indicated as not determined (n.d). D. Humans FLIP-FIGL1 interact in yeast two hybrid assay. Legend as in C.

The distribution of FLIP orthologs in eukaryotic species was analyzed using remote homology search strategy (see Methods). Orthologs of FLIP could be unambiguously detected in a wide range of species including mammalia, sauria and plants but also in arthropods and unicellular species such as choanoflagellate *(Salpingoeca rosetta)* (Figure 2 and as interactive tree http://itol.embl.de/tree/132166555992271498216301). The FLIP orthologs showed low conservation at the sequence level (e.g AtFLIP and hFLIP sharing only 12 % sequence identity), but they all harbor a specific DUF4487 domain (Domain of Unknown Function)^37^, further supporting their orthology. No FLIP ortholog could be detected in alveolata, amoebozoa and fungi. FLIP1 systematically co-occur with FIGL1, which is consistent with FLIP supporting the function of FIGL1 (Figure 2). The reverse is not true since there are a number of species with FIGL1 ortholog detected but no FLIP (as in *D. melanogaster* and *C. elegans*). Structural predictions using RaptorX server^38^. and HHpred ^39^. do not converge toward the same predicted fold but are both in agreement with FLIP likely folding as a long helical bundle over its full sequence. Such folds are often seen in protein recognition scaffolds suggesting FLIP could act as a FIGL1 adaptor module. Given the wide range of species harboring both FLIP and FIGL1 orthologs, the origin of this complex is probably quite ancient at the root of the eukaryotic tree suggesting that absence of FLIP/FIGL1 in eukaryotic species is due to independent gene loss events.

**Figure 2:**
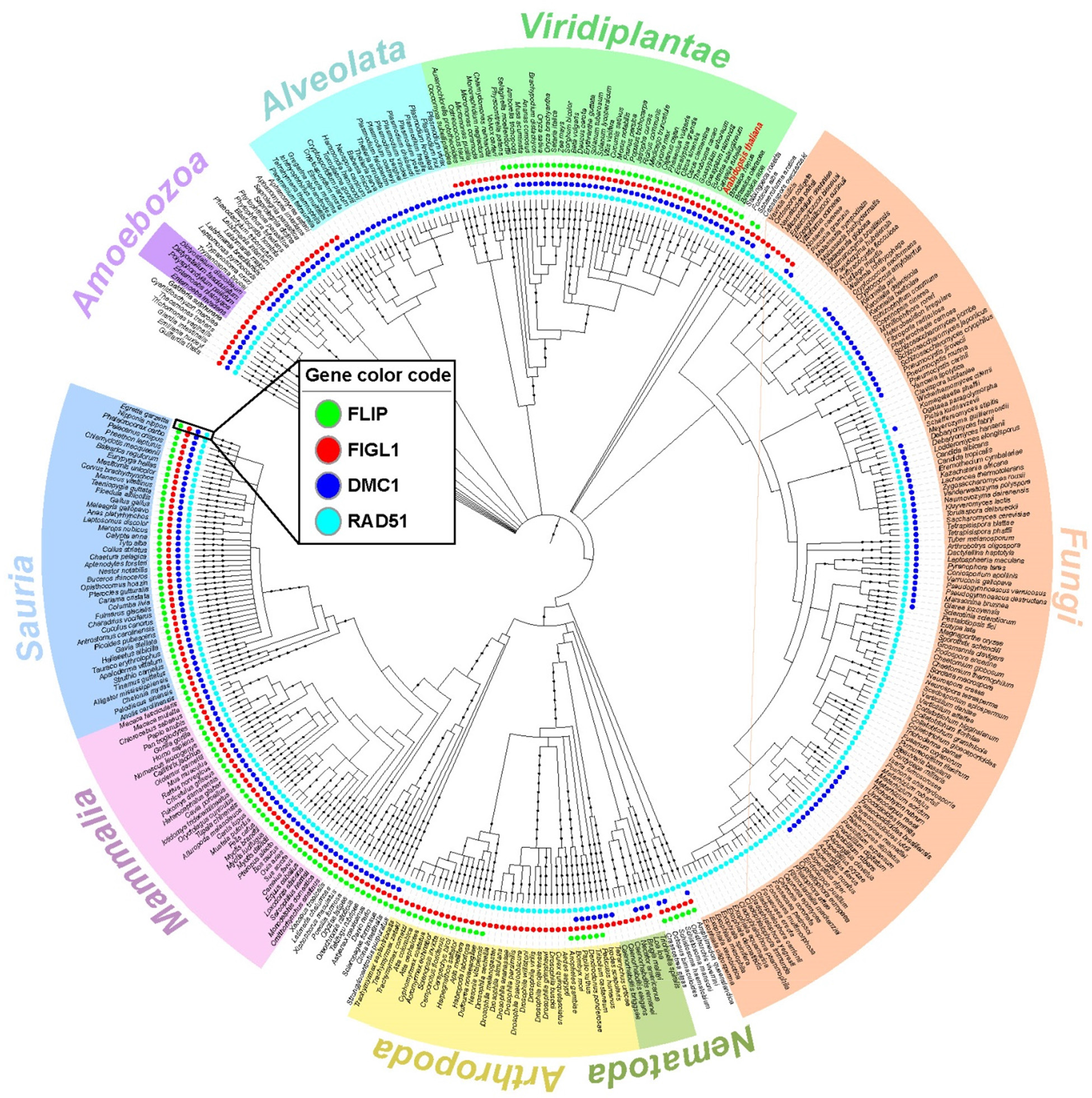
Phylogenetic tree depicting the evolutionary conservation of FLIP, FIGL1, RAD51 and DMC1 orthologs in a range of eukaryotic species. FLIP, FIGL1, DMC1 and RAD51 are presented as dots in green, red, blue and turquoise color, respectively.

### A genetic screen identified FLIP as an anti-CO factor

In parallel, *FLIP* was also recovered in a genetic screen aiming at identifying meiotic anti-CO factors that previously uncovered *FIGL1*. Using fertility (fruit length) as a proxy for CO formation, we screened for ethyl methane sulfonate-generated mutations that restored COs in class I CO deficient mutants (*zmm*). As COs provide a physical link between pairs of chromosomes (bivalents), mutation of an anti-CO factor is expected to restore bivalent formation in CO-deficient mutants, thus improving balanced chromosome segregation and fertility^22^. This genetic screen, led to the identification of several anti-CO factors, defining three pathways that limit COs in Arabidopsis:(i) The FANCM helicase and its cofactors^22,23^; (ii) The AAA-ATPase FIDGETIN-LIKE-1 (FIGL1)^10^; (iii) The RECQ4 helicase-Topoisomerase 3α-RMI1 complex^24,25^. Here, we isolated an additional suppressor of *hei10*, one of the *zmm* mutants that are deficient in class I CO ^40^. This suppressor, *hei10(S)320* showed longer fruit length compared to *hei10* and bivalent formation was restored to an average of 3.7 bivalents per cell compared to 1.5 in *hei10*, and 5 in wild type (Figure 3), suggesting a partial restoration of CO formation. Whole genome sequencing and genetic mapping of *hei10(S)320,* defined a genetic interval containing five putative causal mutations. One of them resulted in a stop codon in the gene *AT1G04650*, which encodes FLIP (*flip-1* W305>STOP). An independent mutation in *FLIP* (T-DNA Salk 037387/ *flip-2*)), was also able to restore bivalent formation in *hei10* (Figure 3). Further, *flip-1/flip-2 hei10* exhibited restored bivalent (Figure 3), demonstrating that *flip-1* and *flip-2* are allelic and that mutations in *FLIP* are causal for the restoration of bivalents in *hei10*. The *flip-1* mutation was also able to restore bivalent formation in *msh5* (Figure 3), another essential gene of the class I CO pathway, suggesting that effect of the *flip* mutation is not specific to *hei10* but allows the formation COs in the absence of the class I pathway.

**Figure 3:**
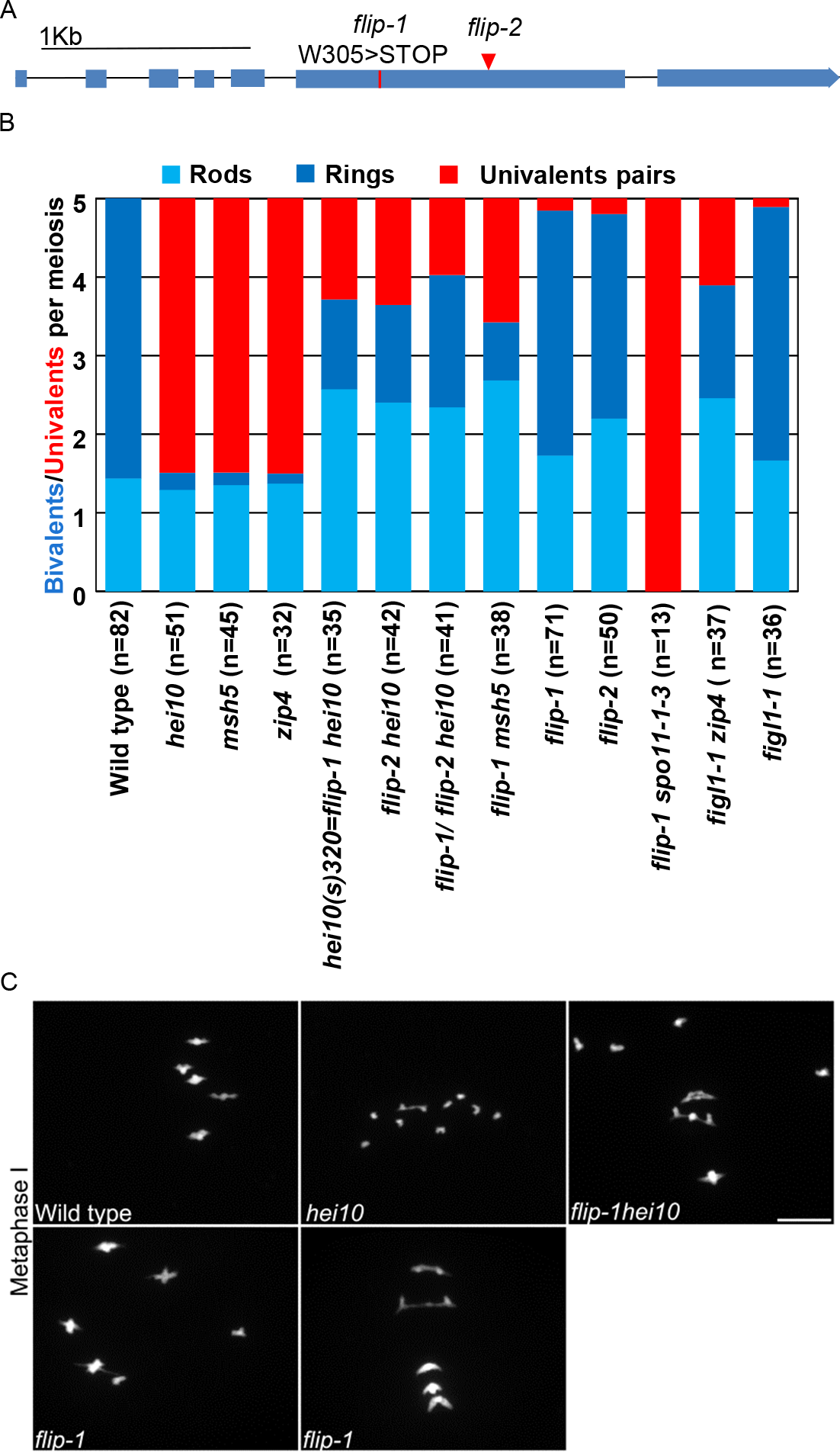
Mutation in *FLIP* restores crossover formation in *zmm* mutants: A. Schematic representation of *FLIP* (Fidgetin-Like-1 Interacting Protein). Exons appear as blue boxes, red line indicates a point mutation and red triangle represents position of T-DNA insertion. B. Average number of bivalents (blue) and pairs of univalents (red) per male meiocytes at metaphase I (Figure C). Light blue represents rod shaped bivalents indicating that one chromosome arm has a at least one CO, and one arm has no CO. Dark blue represents ring shaped bivalent indicating the presence of at least one CO on both chromosome arms. The number of cells analyzed for each genotype is indicated in brackets. Data for *figl1-1, figl1-1 zip4*, and *zip4* was obtained from previous study C. DAPI staining of Chromosome spreads of male meiocytes at metaphase I. Scale bars 10μm.

No growth or development defects were observed in the *flip* mutants. Meiosis progressed normally in single *flip-1* and *flip-2*, except that a pair of univalent was observed at metaphase in 17% of the cells (n=11/71 in *flip-1;* n=9/50 in *flip-2*). (Figure 3B and C). This suggests a slight defect in implementation of the obligate COs in absence of *FLIP*. We next monitored the direct effect of *FLIP* mutation on CO frequency by tetrad analysis and measured recombination in six genetic intervals defined by fluorescent tagged markers that confer fluorescence in pollens ^41^. CO frequency in *flip-1* was significantly increased in four intervals out of six tested, in the range of +15% to +40% compared to wild type (Figure 4). This increase in CO frequency due to loss of *FLIP* is consistent with the restoration of bivalent formation in *zmm* mutants, and implies that FLIP limits COs during meiosis in wild type. FLIP physically interacts with FIGL1 (see above), suggesting that they can act together to limit COs. We therefore compared recombination in *flip-1, figl1-1* and the double mutant by tetrad analysis. On the four intervals tested, *figl1-1* showed an average of ~70% CO increase compared to wild type, corroborating previous findings (Figure 4), which is significantly higher than *flip-1.* Combining *flip-1* and *figl1-1* mutations, did not lead to a further increase in recombination suggesting that *FIGL1* and *FLIP* act in the same pathway to negatively regulate CO formation (Figure 4). However, FIGL1 may be partially active in absence of FLIP as *flip-1* increases CO frequency to a lesser extent than *figl1-1*.

**Figure 4:**
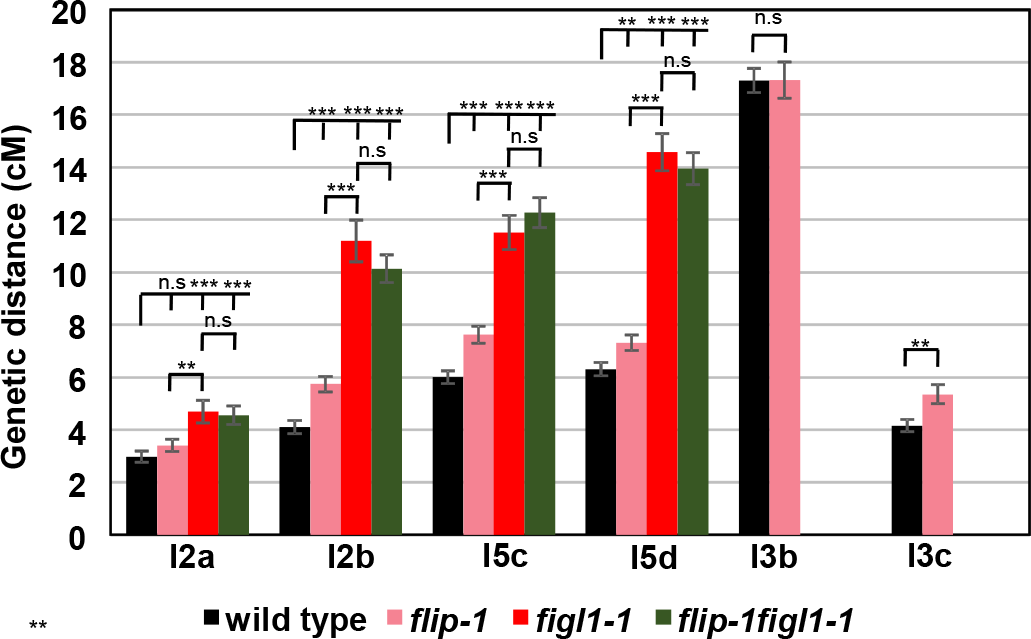
FLIP and FIGL1 act in the same pathway to limit COs. Genetic distance in centiMorgan (cM) measured by pollen tetrad analysis using fluorescent tagged lines ^41^. I2a and I2b are adjacent intervals on chromosome 2. Similarly I3bc and I5cd on chromosome 3 and 5, respectively. Error bar indicates standard error of the mean. Not significant (n.s) *p* > 0.05; ** *p* < 0.01; *** *p* < 0.001, Z-test. Raw data are presented in Table S3.

### *FLIP* limits class II CO

We next explored the origin of extra COs in *flip*. In the *flip spo11-1* double mutant, bivalent were completely abolished and 10 univalents were observed at metaphase I, (Figure 3B), showing that all COs in *flip-1* are dependent on SPO11-1 induced DSBs. Two classes of CO exist in *Arabidopsis*: class I CO are dependent on ZMM proteins and are subjected to interference, while class II are insensitive to interference and involve structure specific endonucleases including MUS81^21^. The *flip-1* mutation restored CO formation in two *zmm* mutants, *hei10* and *msh5* (see above). Further, tetrad analysis of three pairs of intervals showed reduced interference in *flip-1* compared to wild type (Figure 5A). Finally, we examined meiosis in *flip-1 mus81* double mutant. The *flip-1 mus81* double mutant exhibited chromosome fragmentation at anaphase I, which is not observed in either single mutant (Figure 5B). This suggests that MUS81 is required for repair of recombination intermediates formed in *flip-1*. Altogether, the extra COs produced in *flip-1* appeared to be dependent on the Class II pathway, as previously shown for *figl1*.

**Figure 5:**
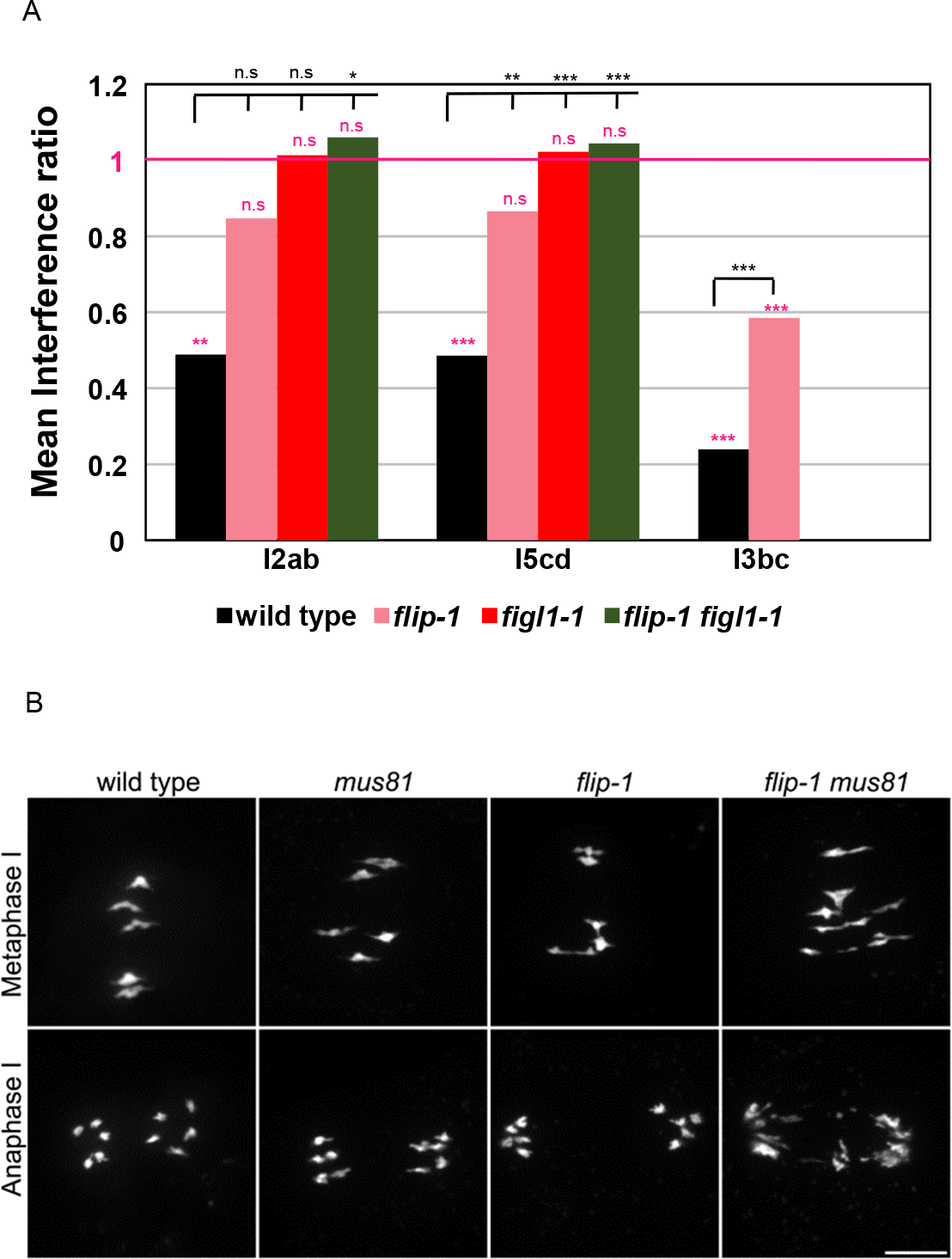
FLIP limits Class II COs. A. Interference ratio is the ratio of the genetic size in an interval with CO in an adjacent interval divided by the genetic size of the same interval without CO in the adjacent interval. This ratio provides an estimate of the strength of CO interference. IR close to 0 means strong interference; Interference ratio =1 (purple line) indicates that interference is absent. The test of absence of interference is shown in purple (n.s *p* > 0.05; ** *p* < 0.01; *** *p* < 0.001). Comparison of Interference data between the genotypes wild type and mutants is indicated in black (n.s *p* > 0.05; * *p* < 0.05 ** *p* < 0.01; *** *p* < 0.001). B. DAPI staining of Chromosome spreads of male meiocytes at metaphase I and anaphase I. Scale bars 10μm.

### Like FIGL1, FLIP negatively regulates DMC1 foci

Based on genetic and physical interactions between FIGL1 and FLIP, we next hypothesized that FLIP might regulate dynamics of DMC1 during meiosis, as previously shown for FIGL1^10^. In wild-type Arabidopsis, DMC1 foci first appear at zygotene and gradually all disappear at pachytene. In contrast, DMC1 foci persist in a fraction of pachytene cells in *flip1-1* (Figure 6A and 6B).This shows that the dynamics of DMC1 foci is modified in absence of *FLIP*, like in absence of *FIGL1*. Persistent DMC1 foci may represent unrepaired DSBs that are eventually repaired, as no chromosome fragmentation was observed at anaphase I (Figure 5B) in *flip-1* mutant.

**Figure 6:**
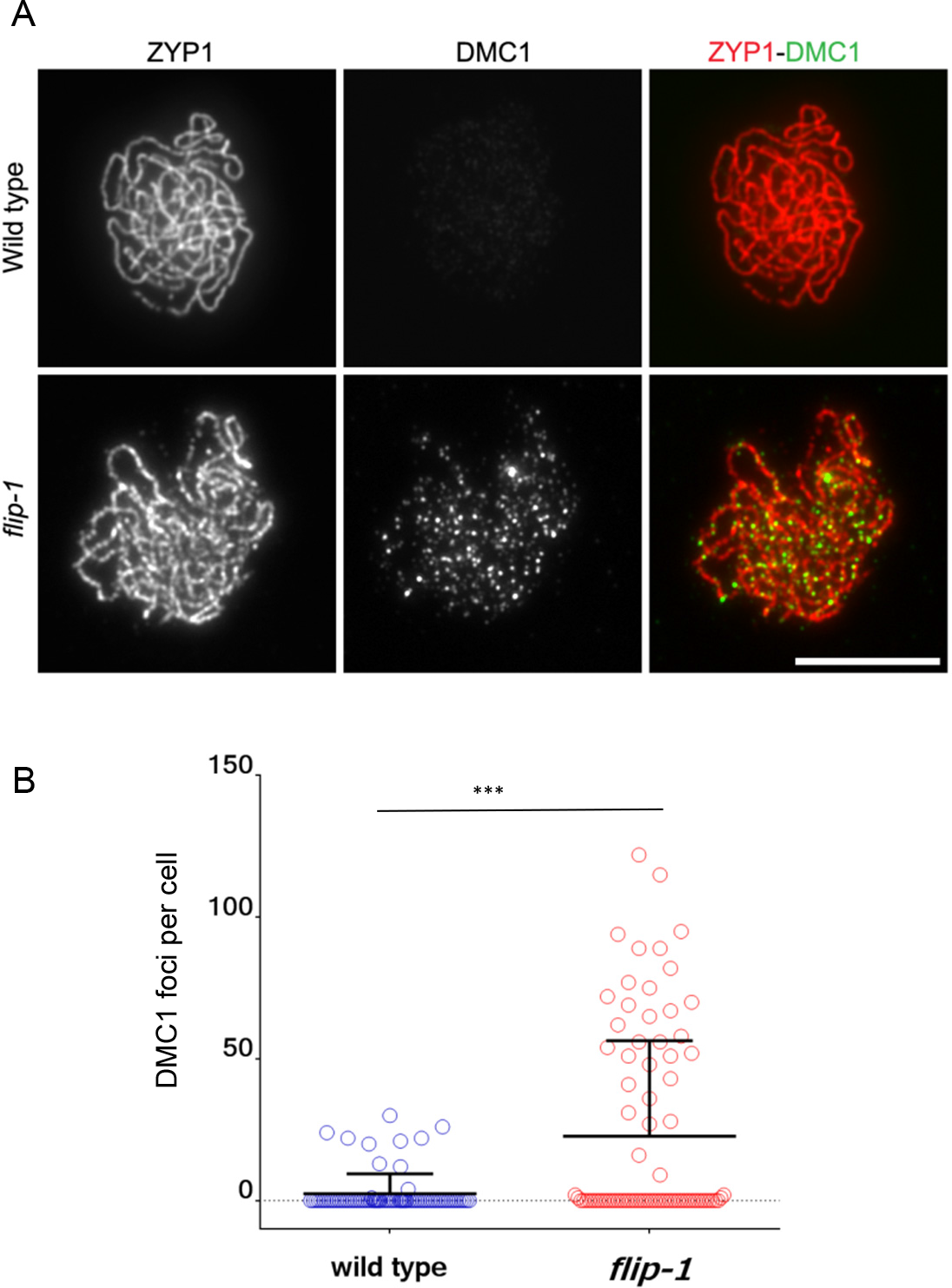
DMC1 foci dynamics is modified in *flip-1*. A. Dual immunolocalization of ZYP1 (red) and DMC1 (green) on meiotic chromosome spreads. ZYP1 marks full synapsis indicative of pachytene stage. Scale bars 10μm. B. Quantification of DMC1 foci per cell in both genotypes. T-test *** *p* < 0.001. Black line indicates mean with standard deviation.

One known positive regulator of DMC1 in plants is SDS, a meiosis-specific cyclin-like protein^42,43^. In absence of SDS, DMC1 foci do not form, synapsis and CO are abolished, but DSBs are formed and repaired presumably using the sister as template ^42,43^. We previously showed that mutation in *FIGL1* restores DMC1 foci formation, synapsis, and bivalent formation in *sds* ^10^. These results argued for antagonistic functions of SDS and FIGL1, the former positively and the later negatively regulating DMC1 foci formation and DMC1-mediated homolog engagement. Here, we similarly showed that DMC1 foci and synapsis are partially restored in *flip-1 sds* double mutant as compared to *sds* (Figure 7A, 7B and 7C). Moreover 4.8 bivalents per metaphase I were observed in *flip-1 sds* (n=10) while their formation is almost completely abolished in *sds* (0.12 bivalents per metaphase I, n=50) (Figure 7D). Taken together, FIGL1 and FLIP could antagonize SDS in the regulation of DMC1 foci formation and DMC1 mediated inter-homolog interactions.

**Figure 7:**
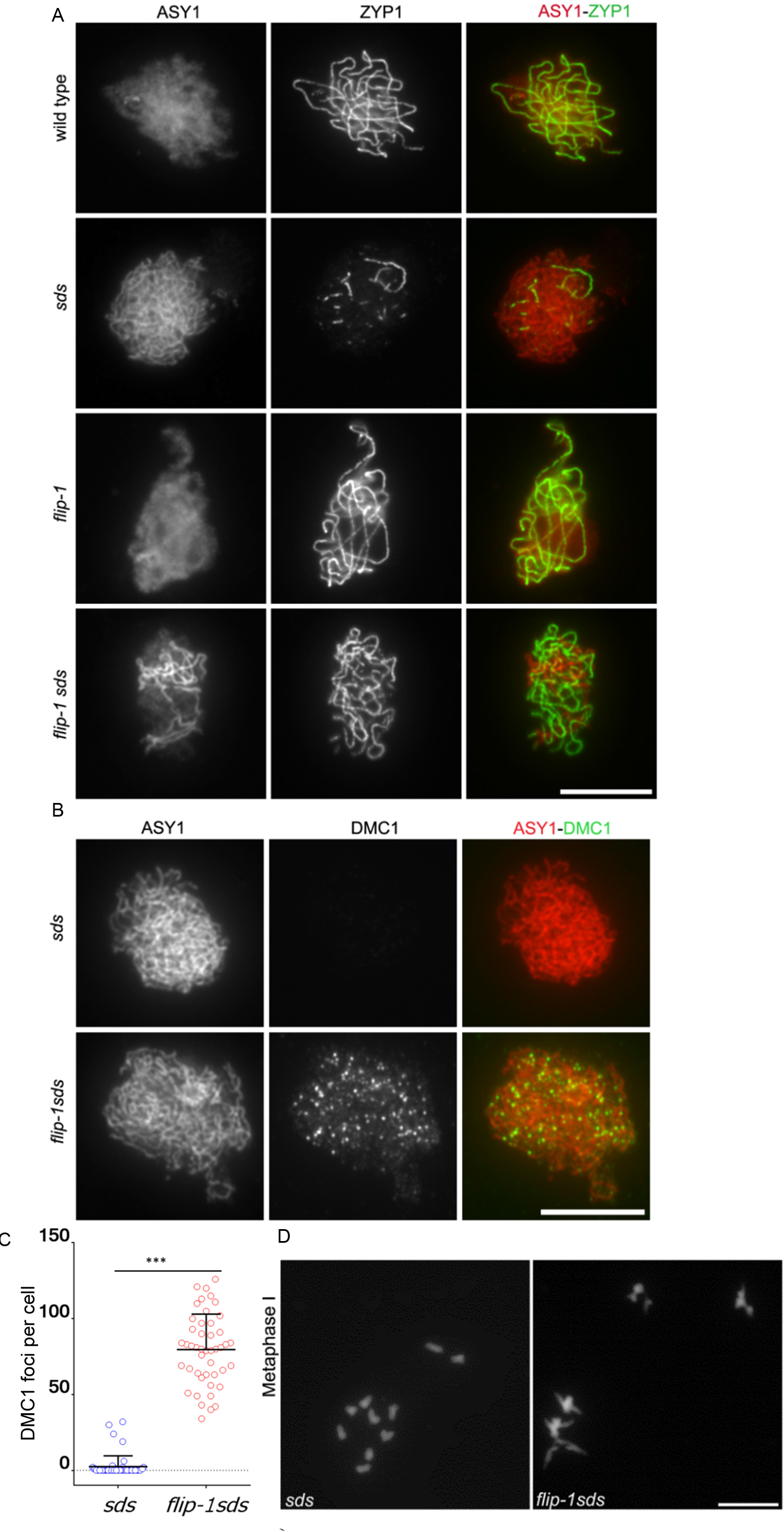
*FLIP* genetically interacts with *SDS*. A. Co-immunolocalization of ZYP1 (green) marks synapsed regions with chromosome axis protein ASY1 (red) on meiotic chromosome spreads. Synapsis was partially restored in *flip-1 sds* compared to single mutant *sds*. Scale bars 10μm. B. Immunostaing of DMC1 (green) and chromosome axis protein ASY1 (red) on meiotic chromosome spreads. C. Quantification of DMC1 foci in *sds* and *flip-1 sds* mutant. T-test *** *p* < 0.001. Black line indicates mean with standard deviation. D. DAPI staining of Chromosome spreads of male meiocytes at metaphase I. Bivalent are restored in *flip-1sds* compared to single *sds*. Scale bars 10μm.

### The FLIP-FIGL1 complex interact with RAD51 and DMC1

Our genetic interaction and immunolocalization studies in Arabidopsis suggest that the FIGL1/FLIP complex might regulate the function of RAD51 and DMC1, directly or indirectly. In addition, it was shown that human FIGNL1 interacts with human RAD51 through its FIGNL1’s RAD51 binding domain (FRBD) ^31^. Hence, we set out to examine whether Arabidopsis and human FIGL1 and FLIP interact with RAD51 and DMC1, using Y2H assays. Consistent with published data, the Y2H assay showed an interaction between the FRBD domain of human FIGNL1 and RAD51 (Figure 8A). Similarly, we detected an interaction between Arabidopsis FIGL1 and RAD51, mediated by the predicted FRBD domain (Figure 8B). In addition, we observed a strong interaction between Arabidopsis FIGL1 and DMC1 as well as between the FRBD domain of the human FIGNL1 and DMC1 (Figure 8). This shows that FIGL1 can interact with both RAD51 and DMC1 and that these interactions are conserved in plants and mammals. Next, we tested interaction between FLIP and the two recombinases, with both plant and human proteins. Human FLIP interacted with DMC1, suggesting that FLIP could reinforce the interaction of the FIGL1-FLIP complex with DMC1. However, our Y2H assay did not reveal any interaction between Arabidopsis FLIP1 and DMC1. Altogether, our data argues that the FIGL1-FLIP complex could directly interact with RAD51 and DMC1.

**Figure 8.**
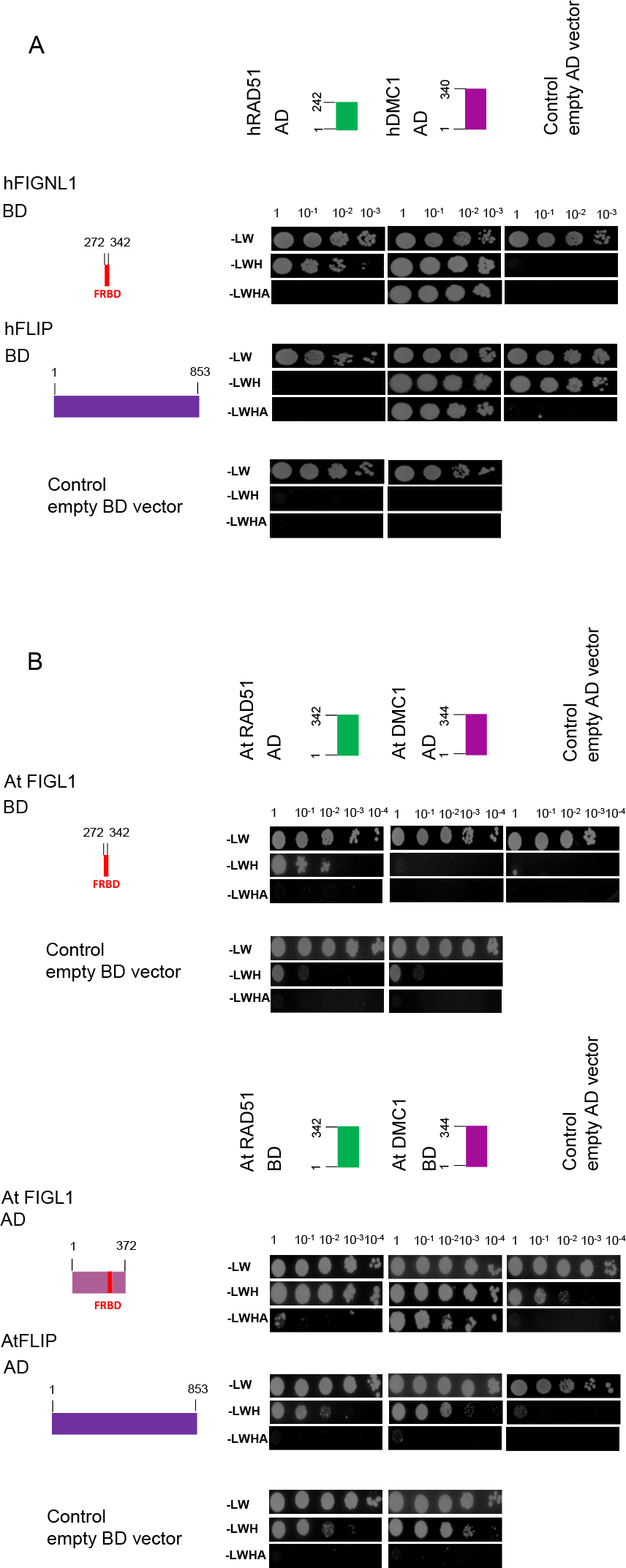
The FIGL1-FLIP complex interact with RAD51 and DMC1 in yeast two hybrid assays. A. The FRBD domain of hFIGNL1 interacts with both human RAD51 and DMC1, while hFLIP was found to interact with only DMC1. Schematic representation of full length and truncated proteins used in this interaction assays. Serial dilutions (1 to 10^−3^) of diploid strains expressing different fusion proteins were tested for interaction by plating on minimal media lacking different amino acids (SD-LW, SD-LWH and SD-LWHA). B. The FRBD domain of Arabidopsis FIGL1 interacts with both RAD51 and DMC1. No interaction of Arabidopsis FLIP and RAD51 as well as DMC1 were detected in yeast two hybrid assays. Full-length and truncated protein used in assay are schematically represented. Serial dilutions of diploids were tested on selection media, as mention above.

## Discussion

Here, we identified by two different approaches FLIP as a new factor that genetically and physically interact with FIGL1^10^ and regulates meiotic recombination. We showed that (i) FIGL1 and FLIP form a conserved complex; (ii) *FLIP* and *FIGL1* are anti-CO factors that act in the same pathway to regulate meiotic recombination; (iii) *FLIP* and *FIGL1* regulate DMC1 foci dynamics; (iv) *flip* and *figl1* restore DMC1 foci formation and DMC1 mediated inter-homolog interactions in the *sds* mutant; (v) FIGL1-FLIP complex interacts with RAD51 and DMC1, and this interaction is evolutionarily conserved in both plants and mammals. FIGL1 was previously shown to be involved in meiotic recombination in *Arabidopsis*, and in recombination-mediated DNA repair in human somatic cells ^31,44^, In contrast and despite the conservation in many eukaryotes, FLIP was of unknown function. With this study, we propose a model wherein FIGL1 and FLIP act as a complex that negatively regulates the strand invasion step of HR by interacting with DMC1/RAD51 and modulating their activity/dynamics (Figure 9). FIGL1 belongs to the AAA-ATPase group of proteins, which typically function by dismantling the native folding of their target proteins ^32,33^. Therefore, it is tempting to suggest that the FLIP/FIGL1 complex may directly disrupt DMC1/RAD51 filaments using the unfoldase activity of FIGL1. Supporting this possibility, both Arabidopsis and human FIGL1 physically interacts with DMC1 and RAD51.

**Figure 9.**
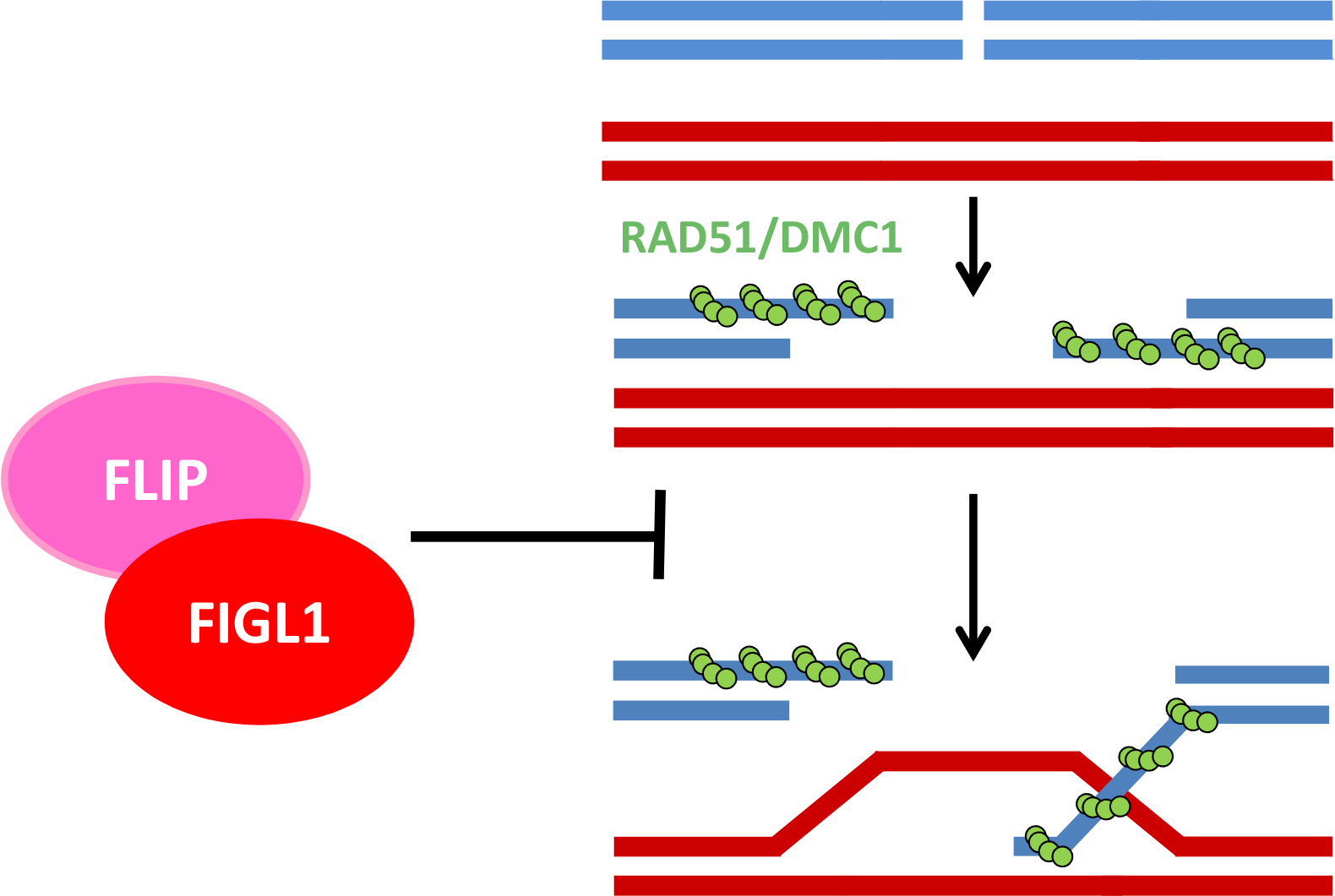
Model: The FLIP-FIGL1 complex controls strand invasion by negatively regulating RAD51/DMC1.

We showed that FLIP1 and FIGL1 act together to limit meiotic CO in Arabidopsis, but the increase in CO frequency is lower in *flip* than in *figl1* (~30% and ~70% increase compared to wild type, respectively). This difference in CO frequency could be attributed to the catalytic activity of the complex being supported by FIGL1. We suggest that FLIP could only be partially required for FIGL1 enzymatic functions *in vivo*, acting as a co-factor or reinforcing the affinity and/or the specificity of the interaction of the FIGL1/FLIP complex with the target. In our assay, human FLIP interacted with DMC1, suggesting that FLIP could indeed function to facilitate FIGL1 activity towards DMC1. We could not detect an interaction between FLIP and RAD51, but we cannot rule out the possibility that FLIP facilitates also interaction of the complex with RAD51. Indeed, several lines of evidence suggest that FLIP could act in conjunction with FIGL1 in its role in somatic HR Down-regulation of *hFLIP* induces reduced growth of HeLa cells *FLIP* in mouse is strongly co-expressed with cancer related genes and the knock out mouse is not viable^36,45^. Finally, *FIGNL1* and *hFLIP* are strongly co-regulated in mouse expression data ^36^. Overall, this argues for a conserved role of the FIGL1/FLIP complex in regulating RAD51/DMC1 activities during both somatic and meiotic HR.

Beyond Arabidopsis and humans, *FIGL1* and *FLIP* are conserved in all vertebrates and land plants we examined. FIGL1 and FLIP can be also detected in species from other distant clades, suggesting that this complex emerged early in the evolution of eukaryotes (Figure 2). However, some clades appear to have lost both *FIGL1* and *FLIP,* most notably the *Alveolata* and *Dikarya* (which regroups the fungi *Basidiomycetes* and *Ascomycetes*). In those species, the RAD51/DMC1 activity might be regulated independently of *FIGL1/FLIP1*. Species with a *FLIP* ortholog also systematically have a *FIGL1*, but the reverse is not true, several species/clades having FIGL1 but no detectable *FLIP orthologs*. This is consistent with our experimental data that argue for FIGL1 being the core activity of the complex and FLIP as a dispensable factor for FIGL1 activity. While *RAD51* appears to be universally conserved, DMC1 is absent in a number of species (Figure 2). Moreover, we could not find any correlation between presence/absence of *FIGL1* or *FLIP* with *DMC1*. Some species have *DMC1* but no *FIGL1/FLIP* (e.g many fungi)), while others have DMC1 and FIGL1 but not FLIP (e;g some nematodes), or *FIGL1* and *FLIP* without *DMC1* (e.g *Chrophyta*). Altogether, our phylogenic analysis supports that neither FIGL1 nor FLIP are specific to DMC1, and that the FIGL1-FLIP complex can regulate the activity of both RAD51 and DMC1. The FIGL1 complex may also have additional functions unrelated to HR^46^.

We suggest that FIGL1 and FLIP could limit strand invasion mediated by RAD51 and DMC1 (Figure 9). How the lack of this function could lead to an increase in the frequency of meiotic CO as observed in *flip* and *figl1?* One possible explanation is that the absence of FLIP and FIGL1 changes the equilibrium between invasions on inter sister versus inter-homolog, leading to the formation of higher numbers of inter homologs joint molecule and eventually more COs. However, DSBs and presumably inter-homologous joint molecules are already in large excess to COs in wild type^21^, making hard to believe that a simple increase in their number would increase CO frequency. Another non-exclusive possibility is that lack of the FLIP / FIGL1 activity generates aberrant recombination intermediates through either multi-invasions or invasion of both ends of a break. The result that the structure specific nuclease MUS81 becomes essential for completion of repair in *figl1* and *flip1*, suggest that indeed some novel class of intermediates arise in these mutants. Thus, we favor the hypothesis that in absence of *FLIP* and *FIGL1* aberrant joint molecules such as multi-chromatid joint molecules ^47,48^ are formed and need structure specific endonuclease to be resolved, leading to increased COs. Therefore, the function of FLIP/FIGL1 in wild type context could prevent formation of aberrant recombination intermediates by functioning as quality control of strand invasion.

In conclusion, we uncovered a conserved FLIP-FIGL1 complex that directly binds to RAD51/DMC1 and could negatively regulate strand invasion during homologous recombination. It would be of particular interest to further study the function of this complex in mammalian systems and in biochemical assays. Unraveling proteins playing a role in HR pathway would provide better understanding related to various inherited diseases in humans pertaining to defects in HR repair proteins ^2^. Targeting HR protein, could increase the sensitivity of cancer cells to anti-cancer drugs ^49^. Thus, FLIP-FIGL1 could represent potential targets for cancer therapy.

## Materials and Methods

<u>Genetic material:</u> The Arabidopsis lines used in this study were: *hei10-2* (N514624) ^40^, *msh5-2* (N526553)^50^, *mus81-2* (N607515) ^18^, *spo11-1-3* (N646172)^51^, *sds-2* (N806294)^42^,*figl1-1* ^10^, *zip4-2* (N568052) ^52^. Tetrad analysis lines (FTLs) used were as follows: I2ab (FTL1506/FTL1524/FTL965/*qrt1-2*), I3bc (FTL1500/FTL3115/FTL1371/*qrt1-2*) and I5cd (FTL1143/FTL1963/FTL2450/*qrt1-2*). FTLs were obtained from Gregory Copenhaver ^41^. Suppressor *hei10(s)320/flip-1* was sequenced using iIlumina technology at the Genome Analysis Centre, Norwich, UK. Mutations were identified through MutDetect pipeline ^23^. The *flip1-1* causal mutation was C to T substitution at the position chr1:1297137 (Col-0 TAIR10 assembly). *flip-2* (N662136) T-DNA mutant was obtained from the Salk collection and distributed by the NASC. The primers used for genotyping are listed in the table S4.

<u>Cytology techniques:</u> Meiotic chromosome from anthers were spread and DAPI stained as previously described ^53^. Immunostaining of male meiotic chromosome spreads was performed as described in Armstrong *et al* ^54^. For Immunofluorescence, primary antibodies were: anti-DMC1 (1:20) ^55^, anti-ZYP1 (1:250) ^56^ and anti-ASY1 (1:250) ^54^. Secondary antibody Alexa fluor®488 (A-11006); Alexa fluor®568 (A-11077); Alexa fluor®568 (A-11075) and super clonal Alexa fluor®488, (A-27034) obtained from Thermo Fisher Scientific were used in 1:250 dilution. Images were obtained using a Zeiss AxioObserver microscope and were analyzed by Zeiss Zen software. In case of DMC1 staining, data was acquired with 1000 ms exposure and DMC1 foci were manually counted. Scatter dot plots and statistical analysis were performed using the software GraphPad Prism 6.

### Recombination measurement

We used FTLs ^41^ to estimate male meiotic recombination rates at six different genetic intervals I2ab, I5cd and I3bc. For each set of experiment heterozygous plants were generated for the pairs of linked fluorescent marker and siblings from the same segregating progeny were used compared recombination frequency between different genotypes. Slides were prepared as described previously ^41^. Tetrads were counted and sorted to specific classes (A to L) ^41^ using a pipeline developed on the Metafer Slide Scanning Platform. For each tetrad, attribution to a specific class was double checked manually. Genetic sizes of each interval was calculated using Perkins equation ^57^ as follows: D=100× (Tetratype frequency+6× Non Parental Ditype frequency)/2 in cM. The Interference ratio (IR) was measured as described previously ^58 41^. Briefly, in two adjacent intervals I1 and I2, genetic size of I1 was calculated for the two populations of tetrads in I2 interval – D1 is at least with one CO in I2; D2 is without CO in I2. The ratio of D1/D2 revealed presence (when IR<1) or absence (when IR is close to 1 or >1) of the interference. A chi square test is performed to test the null hypothesis (H0: D1=D2). The average of the two reciprocals is depicted on the graph (Figure 4A).

### Cloning

Cloning of the *FIGL1* open reading frame (ORF) is described in ^10^. The *AtFLIP* ORF was amplified using gene-specific primer (Table S4) on cDNA prepared from Arabidopsis flower buds (Col-0 accession). The full length or truncated ORFs of *FLIP* were cloned into pDONR207/pDONR201 vectors to produce entry clones. All plasmid inserts were verified by Sanger sequencing. The ORFs for human FIGNL1 (BC051867), RAD51 (BC001459), DMC1 (BC125163) were obtained from the human orfeome collection, while human FLIP (IMAGE clone: 30389801) ORF was ordered from Source BioScience, UK

### Protein-Protein Interaction

#### Yeast two hybrid assay

For yeast two hybrid assays, *AtFIGL1, AtFLIP, AtRAD51 and AtDMC1* as well as their respective human orthologs (*hFignl1, hFlip, hRad51, hDmc1*) were cloned into destination vectors pGBKT7 and pGADT7 by the Gateway technology. The fidelity of coding sequence of all clones was verified by sequencing. Yeast two hybrid assays were carried out using GAL4 based system (Clontech)^59^ by introducing plasmids harboring gene of interest in Yeast strains AH109 and Y187 and interaction were tested as previously described ^60^.

### Tandem Affinity Purification coupled with Mass Spectrometry (TAP-MS)

TAP-MS analysis was performed as described previously^34^. Briefly, the plasmids expressing FLIP or FIGL1 fused to the double affinity GS^rhino^ tag^34^ were transformed into *Arabidopsis* (Ler) cell-suspension cultures. TAP purifications were performed with 200 mg of total protein extract as input and interacting proteins were identified by mass spectrometry using an LTQ Orbitrap Velos mass spectrometer.. Proteins with at least two high-confidence peptides were retained only if reproducible in two experiments. Non-specific proteins were filtered out based on their frequency of occurrence in a large dataset of TAP experiments with many different and unrelated baits as described^34^.

### Bioinformatics

Identification of putative orthologs of FLIP, FIGL1, DMC1 and RAD51 was performed following different strategies based on the sequence divergence and the existence of paralogs. Since FLIP sequence diverged significantly during evolution without detectable paralog, 3 iterations of hhblits^61,62^ against the uniclust30_2017_04 database were sufficient to retrieve 139 sequences belonging to plants and metazoa species. To get NCBI entries of those proteins, a PSSM generated from the recovered alignment was used as input of a jump start PSI-blast ^63^ against the eukaryotic refseq_protein database^64^. For DMC1 and RAD51, reciprocal best hits of blast searches were used to identify the most likely ortholog in every species. First, DMC1 in *H. sapiens* and S. *cerevisiae* sequences were blasted against the refseq_protein database to gather a set of DMC1 candidates. Each of these candidates was reciprocally blasted against the protein sequences of six fully sequenced genomes wherein DMC1 and RAD51 genes could be unambiguously identified and which were chosen spread over the phylogenetic tree (*H. sapiens, S. cerevisiae, C. reinhardtii, T. gondii, P. falciparum, T. cruzi*). Detection of a DMC1 ortholog was considered correct when one of the 6 DMC1 genes was spotted out as best hit with an alignment score at least 10% higher than that of the second best hit, supporting its significantly higher similarity to DMC1 than to RAD51. The same strategy was followed to assign RAD51 orthologs. In the case of FIGL1, large number of paralogs such as spastin, fidgetin, katanin or sap1-like proteins render the global analysis more complex. A phylogenetic tree was initially built focused on the AAA ATPase domain of 600 protein sequences belonging to fidgetin, spastin, katanin, sap1 and VPS4 families. They were aligned using mafft einsi algorithm^65^ and tree was built with PhyML^64^ using the LG model for aminoacid substitution and 4 categories in the discrete gamma model. This prior analysis helped to delineate which homologs could be considered as orthologs of *H. sapiens* and *A. thaliana* FIDGETIN-like proteins. For the 373 fully sequenced species presented in Figure 2, reciprocal blast best hit searches were then performed to retrieve the Fidgetin-like ortholog when present. FIGL1 ortholog candidates were retrieved from a blast of *H. sapiens* and *A. thaliana* FIGL1 sequences against the refseq_protein database and were assessed by reciprocal best hit searches using these candidates as query against genomes of *H. sapiens* and *A. thaliana.* Detection of FIGL1 orthology was assessed if best hit was FIGL1 sequence with an alignment score at least 10% higher than that of the second best hit. For a limited number of species, orthologs were suspected but not identified in any of the NCBI database. Targeted blast searches where then performed on their genomes using the Joint Genome Institute (JGI) server to further probe the existence of these orthologs which could be detected in 7 cases. All the NCBI and JGI gene entries are listed in table S2 and can be easily retrieved from the interactive tree (http://itol.embl.de/tree/132166555992271498216301) ^66^ by passing the mouse over the species names.

## Acknowledgments

We are grateful to Christine Mézard for critical reading of the manuscript. We thank Gregory Copenhaver for providing the FTL lines. This work was funded by the European Research Council Grant ERC 2011 StG 281659 (MeioSight), the Schlumberger foundation for education and research (FSER) and the Simone et Cino del DUCA fundation/Institut de France.

---

Table S1. TAP-MS data

Table S2. NCBI and JGI gene entries of Figure 2.

Table S3. Raw FTL data

Table S4. Genotyping Primers

